# Working memory performance predicts, but does not reduce, cocaine- and cannabinoid-seeking in adult male rats

**DOI:** 10.1101/2024.05.28.596305

**Authors:** Sierra J. Stringfield, Erin K. Kirschmann, Mary M. Torregrossa

## Abstract

**Background:** Cognitive deficits reflecting impaired executive function are commonly associated with psychiatric disorders, including substance use. Cognitive training is proposed to improve treatment outcomes for these disorders by promoting neuroplasticity within the prefrontal cortex, enhancing executive control, and mitigating cognitive decline due to drug use. Additionally, brain derived neurotrophic factor (BDNF) can facilitate plasticity in the prefrontal cortex and reduce drug-seeking behaviors. We investigated whether working memory training could elevate BDNF levels in the prefrontal cortex and if this training would predict or protect against cocaine or cannabinoid seeking.

**Methods:** Adult male rats were trained to perform a ‘simple’ or ‘complex’ version of a delayed-match-to-sample working memory task. Rats then self-administered cocaine or the synthetic cannabinoid WIN55,212-2 and were tested for cued drug-seeking during abstinence. Tissue from the prefrontal cortex and dorsal hippocampus was analyzed for BDNF protein expression.

**Results:** Training on the working memory task enhanced endogenous BDNF protein levels in the prelimbic prefrontal cortex but not the dorsal hippocampus. Working memory training did not impact self-administration of either drug but predicted the extent of WIN self-administration and cocaine seeking during abstinence.

**Conclusions:** These results suggest that working memory training promotes endogenous BDNF but does not alter drug-seeking or drug-taking behavior. However, individual differences in cognitive performance prior to drug exposure may predict vulnerability to future drug use.

## Introduction

Cognitive deficits are often comorbid with substance use disorder. These cognitive deficits reflect a reduction in executive control, the ability of executive systems to regulate behaviors that might contribute to increased substance use. The bidirectional link between cognitive deficits and vulnerability to substance use, in which prior cognitive deficits may increase vulnerability while substance use can cause cognitive dysfunction, indicates a need to investigate mechanisms to improve executive function to encourage resilience and promote positive treatment outcomes (Lees et al., 2021; Melugin et al., 2021; Verdejo-Garcia and Albein-Urios, 2021). Training on cognitive tasks has been proposed as a strategy to counteract the negative effects of substances on executive function by encouraging neuroplasticity that can supplement existing treatment protocols (Wiers et al., 2013; Caetano et al., 2021). For example, cognitive training paradigms that improve working memory, inhibitory control, and goal-directed behaviors have been added to behavioral therapies, yet this training has had limited success in reducing substance use (Cristea et al., 2016; Verdejo-Garcia, 2016; Caetano et al., 2021). One notable outcome from studies of this type is the finding that individual differences in cognitive performance may contribute to substance use outcomes and the efficacy of cognitive training as an adjunctive treatment (Jaeggi et al., 2014). Given the difficulty in disentangling innate versus substance-induced differences in cognitive function in human subjects, there is a need for preclinical models to determine how cognitive training may impact substance use behaviors in the context of innate functional differences (Belin et al., 2016). Moreover, preclinical studies can investigate mechanisms of cognitive training-induced plasticity that might be harnessed for novel treatments.

The prefrontal cortex is required for the maintenance of working memory, which is an active process that allows for short-term storage and manipulation of information (Goldman-Rakic, 1995; Ragozzino et al., 2002; Chai et al., 2018). Working memory deficits are often comorbid with substance use disorders (Albein-Urios et al., 2012; Frazer et al., 2018; Ramey and Regier, 2019), and working memory training has been shown to promote plasticity within the prefrontal cortex (Olesen et al., 2004; Klingberg, 2010; Brooks et al., 2016, 2020; Pappa et al., 2020). Working memory training is one component of cognitive training paradigms that may have a positive impact on behavior, either for treatment of psychiatric disorders where cognitive deficits are likely, or as a generalizable effect in healthy populations (Bickel et al., 2011; Constantinidis and Klingberg, 2016; Brooks et al., 2020; Stavroulaki et al., 2021).

Brain-derived neurotrophic factor (BDNF) is a biological marker that may play a role in working memory training-induced plasticity (Arosio et al., 2021). BDNF is a growth factor that is involved in neuronal development and synaptic plasticity (Park and Poo, 2012; Kowiański et al., 2018). Elevation of BDNF in the cortex may enhance therapeutic approaches to mood disorders, neurodegenerative diseases, and substance abuse by promoting the growth of neural pathways that enhance cognitive control (Bruijniks et al., 2020; Schosser et al., 2022). Elevated BDNF is associated with working memory training, and increasing BDNF expression in the prefrontal cortex has been proposed to promote memory performance for both short-term and working memory (Håkansson et al., 2016; Ney et al., 2020). In addition, BDNF is reduced in the medial prefrontal cortex after self-administration of cocaine and alcohol (Haun et al., 2018; McGinty, 2022), and directly manipulating BDNF in the medial prefrontal cortex of rodents can reduce cocaine and ethanol seeking (Go et al., 2016; Giannotti et al., 2018). Thus, BDNF may be enhanced under certain circumstances, and enhancement with exogenous BDNF can have a beneficial effect on cognitive and substance use behaviors. The effect of naturally increased BDNF on drug-seeking behaviors has yet to be examined.

We chose to investigate the relationship between working memory and the use of two commonly used classes of drugs of abuse: cannabinoids and the stimulant cocaine. We questioned if training on a working memory task itself was sufficient to promote BDNF levels in the medial prefrontal cortex and if that training would then have a protective effect against drug self-administration and relapse-like behavior similar to exogenously administered BDNF. We also questioned if pre-existing deficits in working memory performance would be associated with the extent of drug taking for either drug or the likelihood of cue-induced drug seeking.

## Methods

### Animals

Male Sprague Dawley rats (250-275 g on arrival) were purchased from Envigo (Indianapolis, IN). Animals were housed in a temperature and climate-controlled facility on a 12:12 hour light/dark cycle, all experiments were run during the light cycle. Rats were habituated to the animal facility for at least 3 days before the experiment began. All rats were pair-housed throughout working memory training and single-housed after catheter implantation for the remainder of the experiment. Rats had access to water *ad libitum* and were fed 15 g rat chow per animal immediately after working memory or drug self-administration sessions. All procedures were approved by the University of Pittsburgh Institutional Animal Care and Use Committee and followed the National Institutes of Health Guide for the Care and Use of Laboratory Animals.

### Apparatus

Behavioral training was conducted in standard operant conditioning chambers housed within sound-attenuating cabinets (Med Associates, St. Albans, VT). For working memory training, chambers were configured with a panel containing 5 nose poke apertures on one wall of the chamber, and a house light, sucrose pellet dispenser, and magazine on the opposite wall. During drug self-administration sessions, chambers were configured with two retractable levers with stimulus lights directly above each lever, house light, magazine, and tone generator on the same wall. Rats completed working memory training and drug self-administration in distinct operant chambers.

### Drugs

The synthetic cannabinoid WIN55,212-2 mesylate (WIN, Cayman Chemicals, Ann Arbor, MI) was dissolved in Tween 80 and 0.9% sterile saline. Cocaine hydrochloride was generously supplied by the National Institute on Drug Abuse Drug Supply Program and dissolved in sterile saline to reach a final concentration of 2 mg/ml. All solutions were sterile filtered prior to self-administration.

### Working memory training and home cage controls

Rats were trained on a delayed-match-to-sample working memory task during daily 1-hour sessions (Kirschmann et al., 2017, 2019; Stringfield and Torregrossa, 2021). Rats were trained to perform the task over 6 increasingly difficult stages. In stage 1, rats were trained to respond into any of 5 illuminated apertures to receive a sucrose pellet reward on a fixed-ratio 1 (FR1) schedule of reinforcement. In stage 2, rats learned to respond into a specific illuminated aperture to receive the sucrose pellet reinforcer. In stage 3, rats learned to respond in a single illuminated aperture during a “sample” phase. After a 0.5 s delay, the “choice” phase began where the rat had to respond into the originally sampled aperture from a choice of 3 adjacent illuminated apertures. Once rats met the criteria of 80% correct responses during this stage, the delay between sample and choice phases was increased over the final three stages. Rats experienced 7 delays ranging from 0.5-6 s in stage 4, 0.5-12 s in stage 5, and 0.5-24 s in stage 6. Each delay was presented randomly within a block of trials, and all delays were presented before a new trial block began. Rats remained in stages 2-6 for a minimum of 3 days and performed at least 80% correct trials at the 0.5 s delay before advancing. Rats in the “Simple” group remained in stage 3 throughout training while the “Complex” group advanced to stage 6 of training. Accuracy in stage 3 was used to assign rats to the Simple or Complex group using a balanced design. To equate training duration across groups, rats in the Simple and Complex groups were matched and continued training in stage 3 until their matched counterpart completed stage 6. The average time to complete training was 29 days (range 21-43 days). A separate group of untrained control animals remained in the home cage throughout this period and were weighed and handled daily. Complex and Simple groups were included to assess whether engagement of working memory processes is required for any training-related effects on protein expression or substance seeking behaviors, or if task training in general, independent of working memory, can be effective.

### Surgical procedures

After the working memory training period, all rats were implanted with indwelling catheters for drug self-administration. Catheter implant surgeries were performed as previously described (Bender and Torregrossa, 2021). Indwelling catheters were constructed in house and chronically implanted into the right jugular vein. Rats were anesthetized with 87.5 mg/kg ketamine and 5 mg/kg xylazine. Rats received 5 mg/kg carprofen as an analgesic for 3 days post-operatively. Rats were allowed to recover from surgery for 6-7 days before beginning self-administration sessions.

### Intravenous self-administration and cue-induced drug seeking

WIN and cocaine self-administration were conducted as previously described (Kirschmann et al., 2017; Rich et al., 2019; Bender and Torregrossa, 2021). Rats self-administered 0.0125 mg/kg/infusion WIN on a FR1 schedule of reinforcement. WIN self-administration consisted of daily 1-hour sessions for the first 4 days, followed by 10 days of 2-hour sessions. A separate group of animals self-administered 1 mg/kg/infusion cocaine during daily 1-hr sessions over 14 days. Rats were randomly assigned to left or right active levers and lever assignments were counterbalanced across rats. Responses on the inactive lever were recorded but had no programmed consequences. Pressing the active lever resulted in drug infusion and the presentation of a 10 s compound conditioned stimulus consisting of illumination of the stimulus light above the active lever and presentation of a tone. A 10 s time out followed each reinforced response where the house light was extinguished, and lever presses were recorded but did not result in drug delivery. Catheter patency was tested after the final session by infusion of 0.1 mg of 10 mg/ml methohexital solution, animals without patent catheters were removed from analysis.

Cue-induced drug seeking was measured during 30-minute sessions where the rat was returned to the self-administration chamber and the active and inactive levers were presented. Pressing the active lever resulted in cue presentation, but no drug was delivered.

### Western immunoblot analysis

A separate cohort of rats that completed working memory behavioral training or remained in the home cage throughout the training period was euthanized by rapid decapitation 24 hours after a final training session. Brains were rapidly frozen in isopentane cooled with dry ice and stored at -80 °C before additional processing. Tissue punches from the prelimbic prefrontal cortex and dorsal hippocampus were collected from 1 mm-thick coronal sections. Tissue was fractioned into membrane-bound and soluble components as previously described (Bañuelos et al., 2014; Kirschmann et al., 2017; Stringfield and Torregrossa, 2021). Protein concentrations were assessed using a bicinchoninic acid assay (Thermo-Scientific Pierce, Waltham, MA). 20 µg protein was loaded into 4-20% Tris-glycine gels (Invitrogen, Carlsbad, CA) for separation by SDS-PAGE. Proteins were transferred to polyvinylidene fluoride (PVDF) membranes and blocked for 1-hour in 5% nonfat dry milk in blocking solution containing PBS and 0.1% Tween 20 (PBST). All antibodies were diluted in 1:1 PBST and Li-COR Odyssey blocking buffer (Li-Cor, Lincoln NE). Membranes were incubated in primary antibody overnight at 4°C, washed, and incubated in secondary antibody for 1 hour at room temperature. Membranes were imaged using the Li-Cor Odyssey image system and analyzed using Li-Cor Image Studio software. Quantified protein bands were normalized to GAPDH loading control and within-gel home cage control samples. Primary antibodies were BDNF (1:1000, Abcam, Cambridge, UK); and GAPDH (1:1000, MilliporeSigma). Secondary antibodies were (IRDye 800CW anti-rabbit, 1:5000, and IRDye 680 CW anti mouse, 1:5000).

### Statistical analysis

Statistical analysis was completed using GraphPad Prism software version 9.2 (GraphPad, San Diego, CA). Performance on the working memory task was calculated by dividing the number of correct trials by the total number of trials completed at each delay length. Accuracy on the working memory task was analyzed by 2-way repeated measures ANOVA (α=0.05 for all analyses) with Bonferroni corrected multiple comparisons completed when applicable. Group differences throughout drug self-administration and across cued drug-seeking tests were analyzed by 2-way repeated measures ANOVA. Protein expression from western blot analysis was compared across groups by 1-way ANOVA. Correlations between working memory performance and drug intake were completed using simple linear regression.

## Results

### Working memory training impacts on BDNF protein concentration

Given that one of our hypotheses was that working memory training might influence prelimbic prefrontal cortex (PrL) function in a manner protective against subsequent substance use related behaviors, we first assessed expression of a plasticity related protein in tissue from rats that completed the “Simple” working memory task, “Complex” task, and Untrained controls by western blot (n=10-12 per group). BDNF protein in the PrL was significantly increased in rats that completed both Simple and Complex training compared to untrained animals (F _(2, 30)_ = 8.78, p=0.001, Fig 1*a*) but not in the dorsal hippocampus (DH) (F _(2, 30)_ = 0.70, p=0.50 Fig 1*b*), which was used as a comparator region. These data suggest that training male rats on both simple and complex operant tasks prominently increases BDNF expression specifically in the PrL.

**Figure 1:**
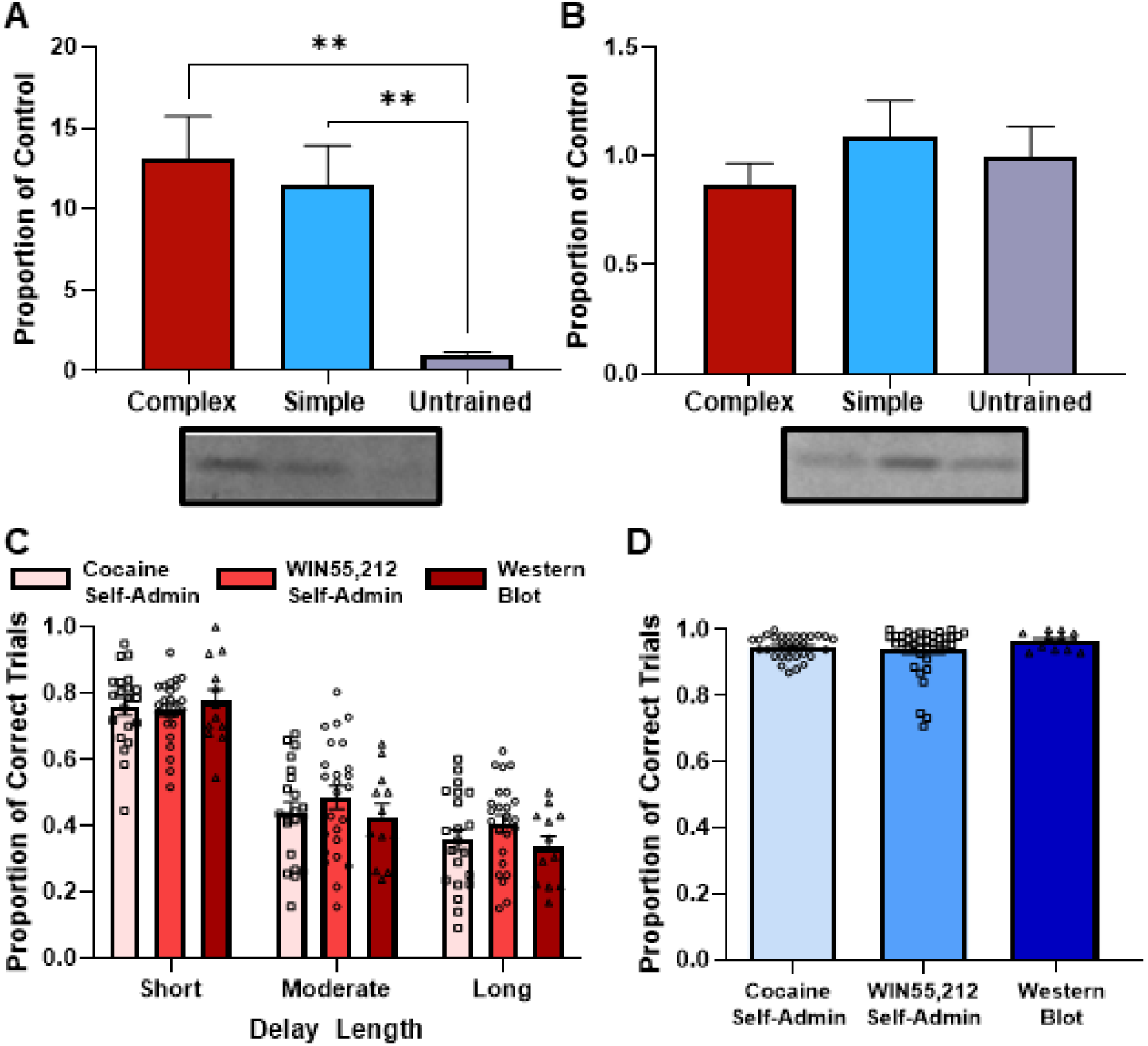
BDNF protein expression after working memory training. Protein concentration from western blot analysis of BDNF in (A) the prelimbic prefrontal cortex and (B) the dorsal hippocampus after training on the Complex and Simple working memory task compared to Untrained control animals (n = 10-12 per group). Protein concentration is expressed as a proportion of Untrained control expression. ** p<0.01. (C) Performance of animals in the Complex group on the working memory task on short (0-4 s), moderate (8-16 s) or long (20-24 s) delays prior to cocaine or WIN self-administration or tissue collection for western blot. (D) Performance of rats in the Simple groups prior to drug self-administration or tissue collection for western blot.

### Effects of working memory training on cocaine self-administration and drug-seeking in abstinence

All cohorts of rats in the Complex and Simple groups were compared for performance on the working memory task. No differences in performance emerged for any measures across cohorts of animals trained on the Complex task that would progress to cocaine or WIN self-administration, or inclusion in western blots (Fig 1*c*, no main effect of group, F _(2, 55)_ = 0.71, p=0.49, no group by delay interaction, F _(4, 110)_ = 0.98). A main effect of delay did emerge across all groups, as performance decreased with increasing delay length (F _(2, 110)_ = 170.8, p<0.001). Similarly, no significant difference emerged across cohorts of animals in the Simple group (Fig 1*d*, F _(2, 77)_ = 1.06, p=0.35).

Based on prior research suggesting that PrL BDNF reduces multiple forms of cocaine-seeking behaviors (Go et al., 2016; McGinty, 2022), we next investigated whether working memory training would influence cocaine self-administration or cue-induced cocaine seeking over the course of abstinence. Rats completed Simple (n=31) or Complex (n=21) working memory training or remained in the home cage (n=21) throughout the same period. These rats self-administered 1 mg/kg/infusion cocaine during daily 1-hour sessions for 14 days (Fig 2). No differences emerged between groups for the number of infusions received (Fig 2*a*), where animals increased cocaine intake over time as represented by a main effect of day (F _(13,91)_ = 18.90, p<0.0001). There was a main effect of group (F _(2, 70)_ = 3.71 p=0.029), and a day by group interaction (F _(26, 910)_ = 2.00, p=0.002) where animals In the Simple and Complex training group received more infusions than Untrained animals over the first 6 days of training, which we attribute to the fact that the rats were experienced with making responses in operant boxes for sucrose pellets rewards due to cognitive task training, and thus they acquired lever pressing more quickly than the untrained group. There was no main effect of group for active lever presses (Fig 2*b*, F _(2, 70)_ = 1.76, p=0.18), although there was a main effect of day as the animals increased their lever pressing as they acquired the task (F _(13,91)_ = 5.93, p<0.0001) as well as a group by day interaction (F _(26, 910)_ = 1.66, p=0.020) where animals in the Complex and Simple tasks pressed more than untrained animals on the second and third days of acquisition. Inactive lever presses (Fig 2*c*) did not differ over time as shown by a lack of a main effect of day (F _(13,91)_ = 1.684, p=0.16), group (F _(2, 70)_ = 0.6371, p=0.41), or day by group interaction (F _(26, 910)_ = 0.9640, p=0.94) for all analyses.

**Figure 2:**
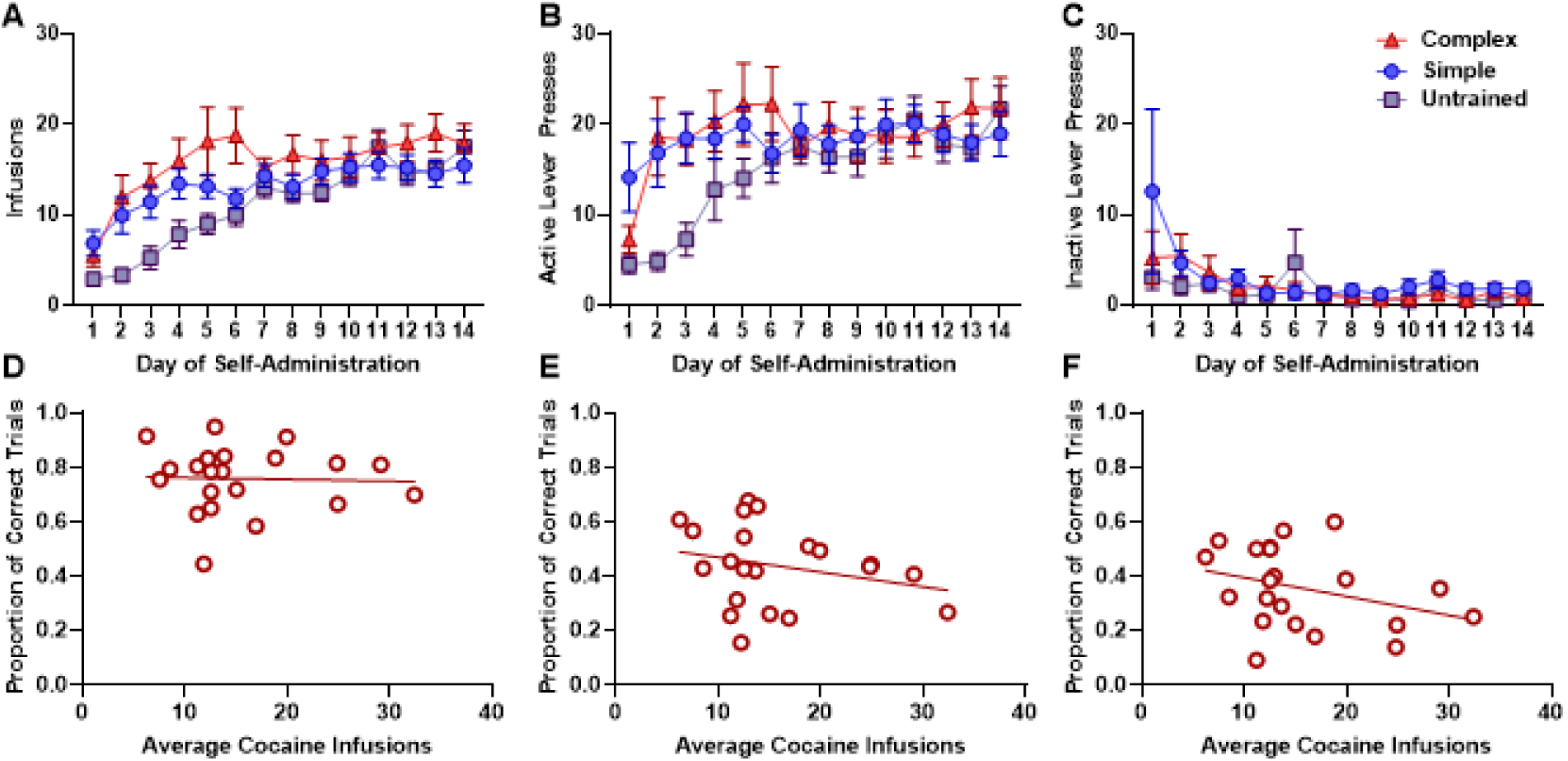
Cocaine self-administration after working memory training. Rats in the Complex (n=21), Simple (n=31), or Untrained (n=21) groups self-administered cocaine (1 mg/kg/infusion) for 14 days. Mean (± SEM) (A) Infusions, (B) active lever responses, and (C) inactive lever responses for all groups throughout cocaine self-administration. (D-F) Correlations between the proportion of correct trials completed on the last day of training on the Complex working memory task and the average number of cocaine infusions received during self-administration. (D) Short delay lengths, (E) moderate delay lengths, (F) long delay lengths.

The relationship between Complex working memory training and cocaine self-administration was further evaluated by correlating the average number of cocaine infusions received with performance on the final day of working memory training (Fig 2*d-f*). No correlation between working memory performance and cocaine intake emerged for short (Figure 2*d*, R^2^=0.001, p=0.87), moderate (Fig 2*e*, R^2^=0.064, p=0.24), or long (Fig 2*f*, R^2^=0.11, p=0.14) delays.

Next, rats that completed Simple or Complex working memory training and untrained controls were evaluated for cued cocaine-seeking (Fig 3). We tested animals for drug-seeking on days 1,7, and 35 of abstinence. A main effect of day emerged for active (Fig 3*a*, F _(2, 98)_ = 25.61 p<0.001) and inactive (Fig 3*b*, F _(2, 98)_ = 4.74 p=0.011) lever presses, but rats reduced lever presses over time instead of demonstrating an increase in responding over increasing abstinence. No main effects of group or group by day interactions emerged for active or inactive presses (p>0.1 for all analyses).

**Figure 3:**
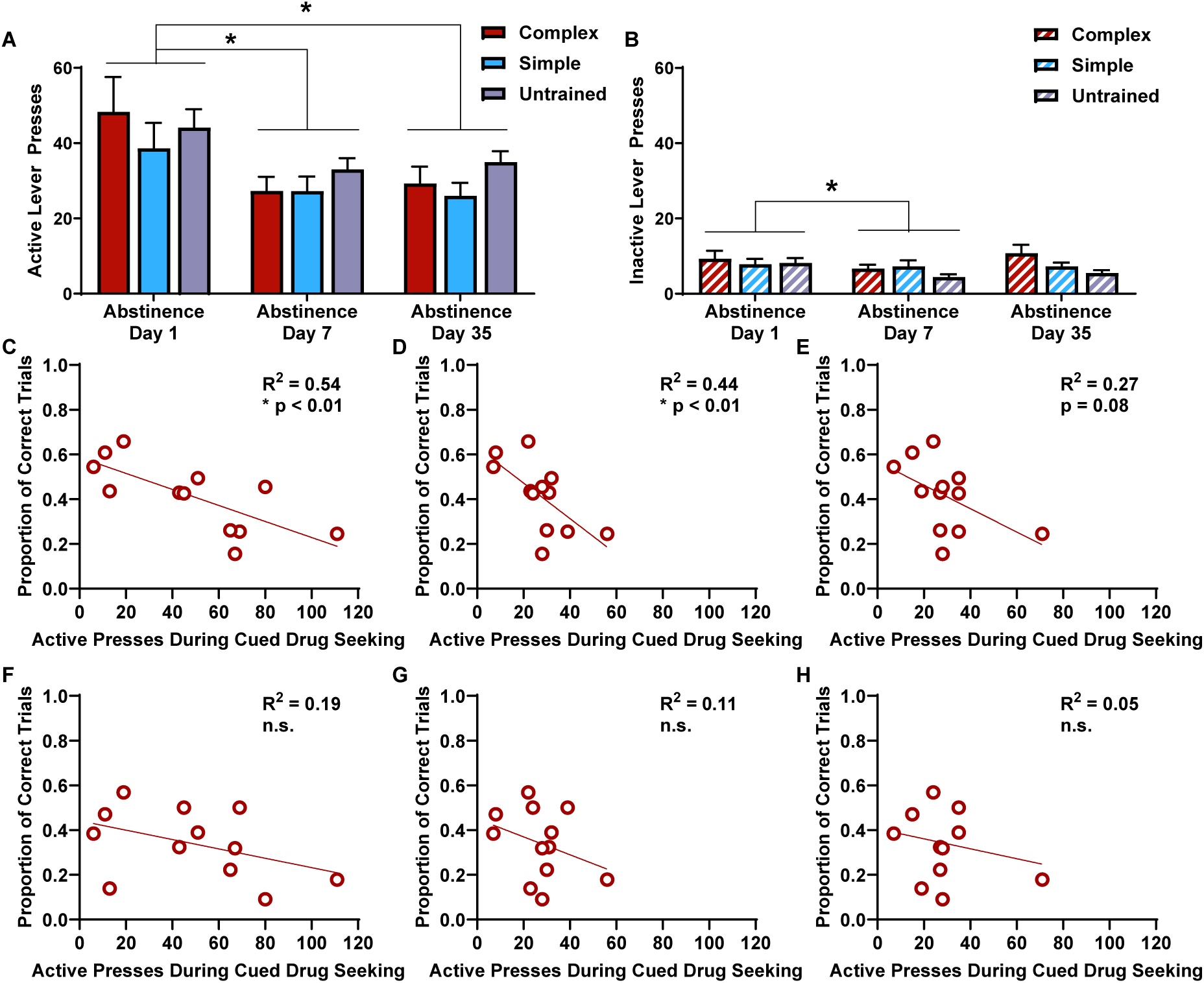
Relationship between working memory training and cocaine-seeking in abstinence. Rats in the Complex (n=21), Simple (n=31), and Untrained (n=21) groups were tested for cued cocaine seeking on abstinence days 1, 7, and 35. (A) Mean (± SEM) Active and (B) Inactive lever responses on each day of testing. (C-E) Proportion of correct trials on the last day of working memory training for moderate delay lengths correlated with active lever presses during testing on (C) Day 1, (D) Day 7, and (E) Day 35 of abstinence. (F-H) Proportion of correct trials completed at long delay lengths correlated with active lever presses during testing on (F) abstinence Day 1, (G) Day 7, and (H) Day 35. * p<0.05.

Relationships between performance on the Complex working memory task and cue-induced cocaine-seeking were evaluated by correlating the proportion of correct trials completed on the last day of training with active lever presses during each day of cued cocaine-seeking tests (Fig 3*c-h*). No significant correlations emerged for short delays (p>0.1 for all analyses) but significant negative correlations emerged at moderate delay lengths on abstinence day 1 (Fig 3*c*, R^2^=0.54, p=0.006) and day 7 (Fig 3*d*, R^2^=0.44, p=0.019) with a trend on day 35 (Fig 3*e*, R^2^=0.27, p=0.08) such that better working memory performance corresponded to fewer active lever presses. No significant correlations emerged at the longest delays on any day of abstinence, though the slopes look similar to the correlations with performance at the moderate delays (Fig 3*f-h*, p>0.1 for all analyses). Overall, these results suggest that while cognitive training *per se* is not protective against cocaine-motivated behaviors, trait level differences in working memory performance may affect individual propensity to exhibit cue-induced cocaine seeking.

### Effects of working memory training on cannabinoid self-administration and drug seeking in abstinence

Given that cognitive training had little effect on cocaine seeking, we wanted to investigate if there would be an effect on a different class of drug, particularly one that is thought to negatively impact working memory, so we next investigated effects on cannabinoid seeking behavior. We used self-administration of the cannabinoid receptor agonist WIN55,212-2 as it is more readily self-administered by rats than THC (Fattore et al., 2001; Lefever et al., 2014). A total of 71 animals (n=37 Simple, n=25 Complex, n=9 Untrained) successfully completed self-administration of 0.0125 mg/kg/infusion WIN. WIN self-administration over 14 days of training was compared for rats that completed the Simple or Complex versions of the working memory task and untrained animals that remained in the home cage during that time (Fig 4). During self-administration, a main effect of day (F _(26, 884)_ = 1.88, p=0.005) and a day by group interaction (F _(13, 884)_ = 2.92, p<0.001) emerged for the number of infusions (Fig 4*a*). The same was present for responses on the active lever (Fig 4*b*), where there was a main effect of day of self-administration (F _(13, 884)_ = 2.48, p=0.003) and a day by group interaction (F _(26, 884)_ = 1.74 p=0.013). A main effect of day was also present for inactive lever presses (Fig 4*c*, F _(13, 884)_ = 1.92, p=0.024). No differences emerged between rats that received training on the Simple or Complex behavioral task for active lever presses, inactive lever presses, or the number of infusions received. While rats that had received prior behavioral training on the Simple or Complex tasks exhibited more active lever pressing and received more infusions during the first half of self-administration, no difference remained between groups after acquiring the behavior. This is consistent with our cocaine self-administration data, indicating that prior operant training can promote faster acquisition of drug self-administration but suggests that working memory training is not protective against acquisition of cannabinoid self-administration.

**Figure 4:**
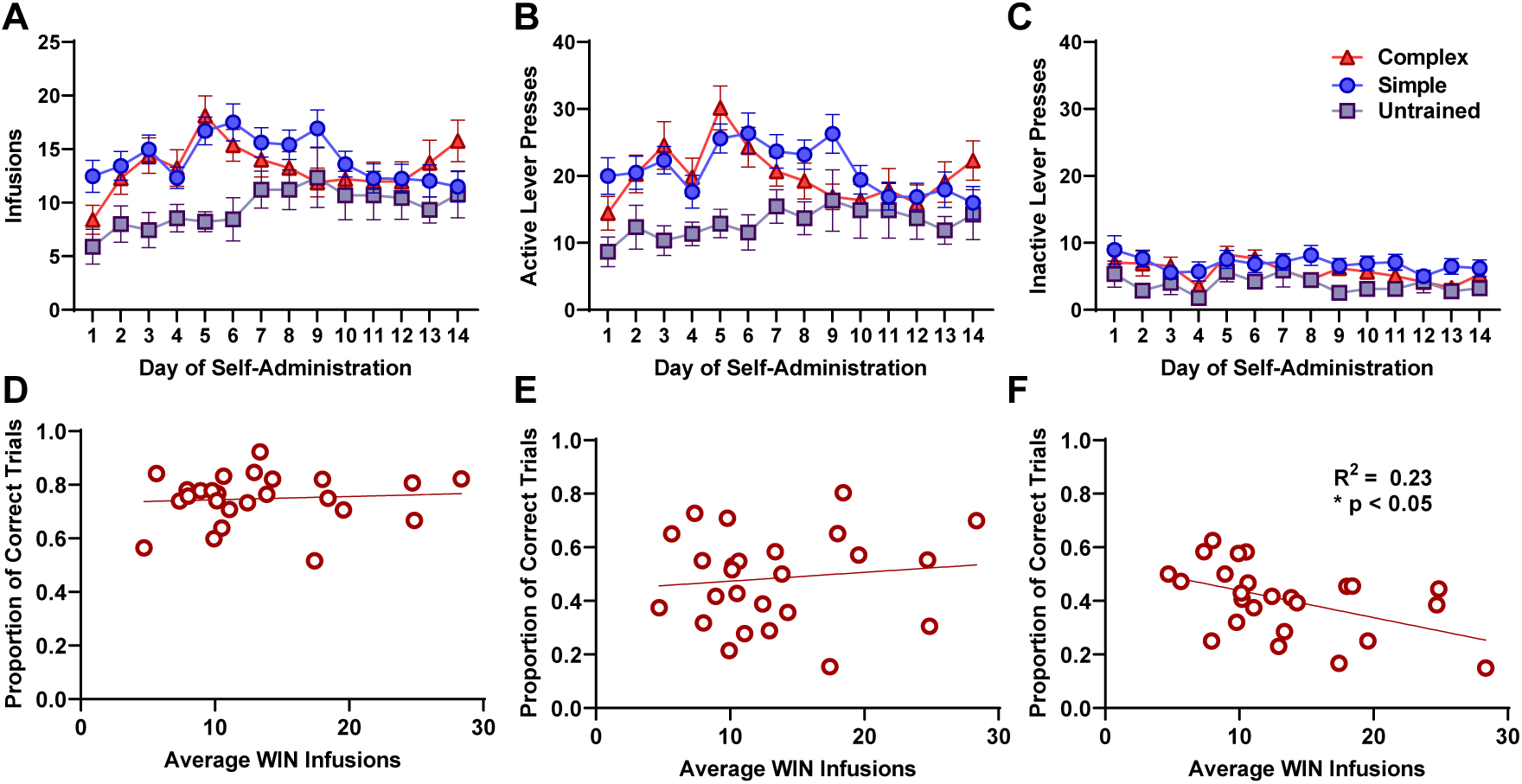
WIN55,212-2 self-administration after working memory training. Self-administration of WIN over 14 days of training. Mean (± SEM) (A) infusions, (B) active lever responses (C) inactive lever responses on each day of self-administration for rats from the Complex (n=25), Simple (n=37), and Untrained (n=9) groups. (D-E) Performance on the Complex version of the working memory task correlated with the average number of WIN infusions completed during self-administration for (D) short, (E) moderate, and (F) long delay lengths. *p<0.05.

To evaluate the extent that innate differences in working memory performance on the Complex task correlated with self-administration of WIN, we compared rats’ performance on the final day of working memory training on short, moderate, and long delay lengths with their average WIN intake during self-administration (Fig 4*d-f*). Accuracy was not associated with WIN intake during self-administration on short (Fig 4*d*, R^2^=0.0066, p=0.7) or moderate length delays (Fig 4E, R^2^=0.014, p=0.57) while performing the Complex task. However, low accuracy at the longest delays correlated with increased WIN intake (Fig 2*f*, R^2^=0.23, p=0.015), where rats that performed worse on these difficult trials would proceed to self-administer more WIN. Thus, these data suggest modest support for the hypothesis that increased innate working memory performance is protective against cannabinoid use-related behavior.

To compare the effects of different degrees of working memory training on drug-seeking in abstinence, a subset of rats trained on the Simple (n= 27) and Complex (n=25) tasks were tested for cued WIN-seeking at three timepoints during abstinence (Fig 5). Tests took place on days 7, 21, and 35 or 36 of abstinence. There were no significant differences between training groups on any day of abstinence for responding on the active lever (Fig 5*a*), nor main effects of working memory training (F _(1, 50)_ = 1.00, p=0.32), day of abstinence (F _(2, 100)_ = 1.293, p=0.28), or a day by group interaction (F _(2, 100)_ = 0.006, p=0.99). An interaction between group and day did emerge for inactive lever presses (Fig 5*b*), where rats trained on the simple working memory task responded more than those trained on the Complex task only on abstinence day 7 (F _(2, 100)_ = 3.84, p=0.025). These results indicate that the complexity of working memory training did not impact drug-seeking in abstinence.

**Figure 5:**
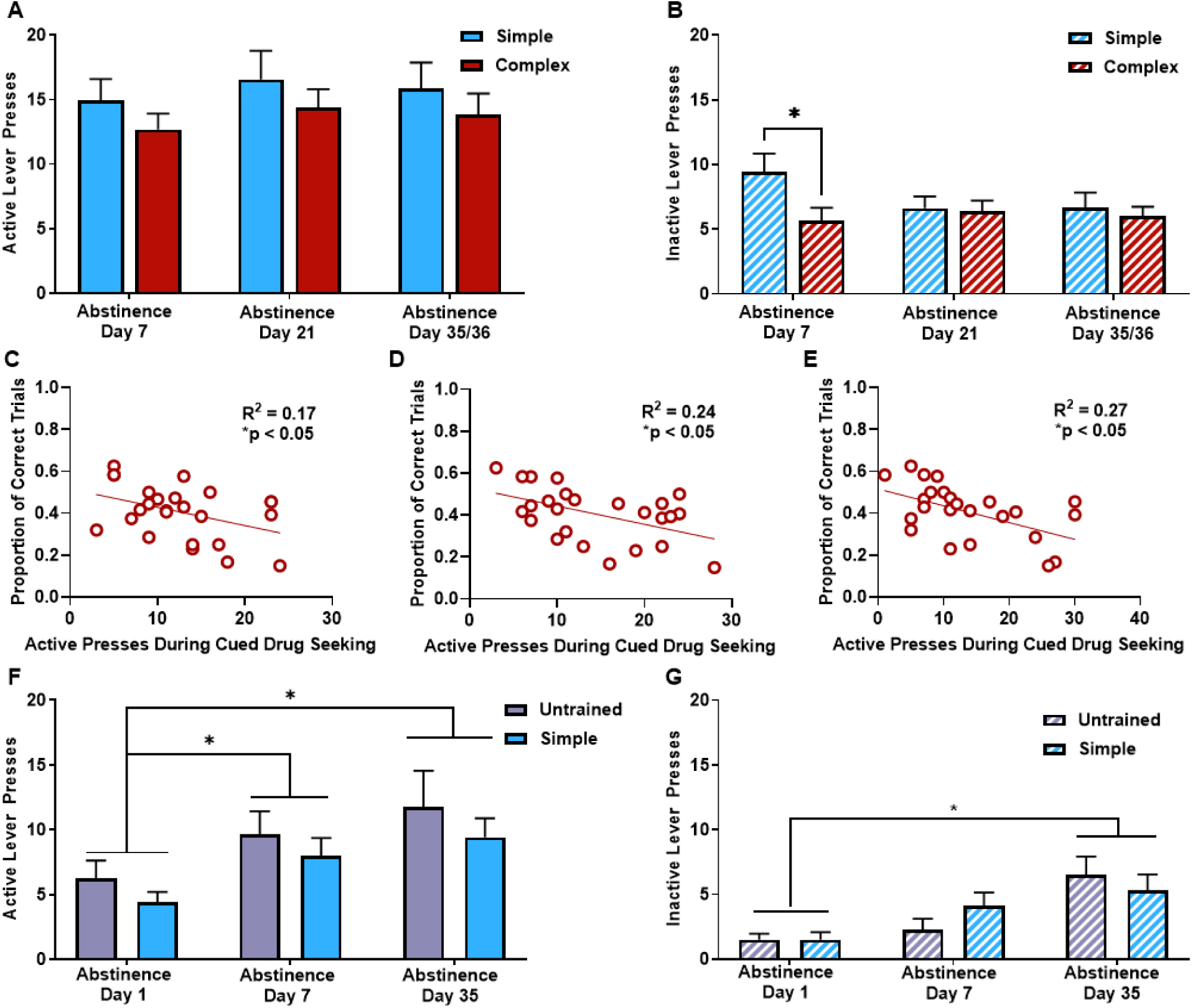
Relationship between working memory training and WIN-seeking in abstinence. Rats trained on the Complex (n=25) and Simple (n=27) working memory tasks were compared for cued WIN-seeking on abstinence days 7, 21, and 35. (A) Active lever responses and (B) inactive lever responses. (C-E) Correlations between the proportion of correct trials completed at long delays on the Complex working memory task on (C) abstinence Day 7, (D) Day 21, and (E) Day 35. (F-G) Rats trained on the Simple working memory task (n=10) and Untrained controls (n=9) were compared for WIN-seeking on abstinence days 1, 7, and 35. (F) Responses on the active lever and (G) responses on the inactive lever. *p<0.05.

As poor performance on the Complex task at long delays was correlated with increased WIN self-administration, we investigated if this correlation emerged for WIN-seeking across abstinence. The number of active lever presses completed on each day of cued WIN-seeking was negatively correlated with the proportion of correct trials completed on the last day of training on the Complex task. Performance on the short and moderate length delays was not correlated with lever responses (p>0.1 for all analyses), but low performance on the longest delays was correlated with increased drug-seeking during cued seeking tests on day 7 (Fig 5*c*, R^2^=0.17, p=0.05), day 21 (Fig 5*d*, R^2^=0.24, p=.014), and day 35/36 (Fig 5*e*, R^2^=0.27, p=0.008). Thus, these data are consistent with the self-administration results above and the cocaine-seeking data, suggesting that better working memory performance may be protective against relapse-like behavior for multiple classes of substances.

Given that the complexity of working memory training did not significantly impact cued drug-seeking in abstinence, we next asked if any behavioral training, even on the simple task, would result in reduced drug-seeking compared to untrained animals. We also questioned if there would be any increase in drug-seeking throughout abstinence, similar to the incubation of craving effect seen with other drugs of abuse including what is traditionally observed for cocaine and alcohol. A separate cohort of rats was trained on the simple version of the working memory task (n=10), while a group of untrained animals remained in the home cage during the same period (n=9). These rats completed WIN self-administration as described above and were tested for cued WIN-seeking (i.e., incubation of craving) on days 1, 7, and 35 of abstinence (Fig 5*f-g*). Behavioral training did not reduce drug-seeking in abstinence compared to untrained animals (F _(1, 17)_ = 1.191, p=0.29), nor was there an interaction between behavioral training and day of abstinence (F _(2, 34)_ = 0.04062 p=0.96). There was a main effect of day, with cued WIN-seeking increasing throughout abstinence compared to day 1 (Fig 5*f*, F _(2, 34)_ = 8.399, p=0.001). Inactive lever pressing from each group followed the same pattern with no significant effect of group or day by group interaction, but a main effect of day emerged indicating an increase in inactive lever responding over time (Fig 5*g*, F _(2, 34)_ = 14.35, p<0.001). Again, these data are consistent with the conclusion that prior cognitive training is not in and of itself protective against incubation of craving, but rather implicates that innate working memory performance is protective against some drug seeking behaviors.

## Discussion

In this study, we investigated the link between working memory performance and drug-seeking behaviors. We found that training on a working memory task, regardless of the complexity, promoted expression of BDNF compared to untrained animals. The change in BDNF was particularly relevant in the prelimbic prefrontal cortex, where behavioral training markedly increased BDNF expression. Next, we found that adult male rats trained on either version of the working memory task or untrained controls showed similar self-administration of the synthetic cannabinoid WIN55,212-2 or cocaine. These animals also did not differ significantly in the extent of cue-induced drug-seeking measured across multiple days of abstinence, indicating that although working memory training promoted BDNF expression in the prefrontal cortex, it did not influence drug-seeking. We compared working memory performance in individuals trained on the complex version of the task with the extent of drug taking and drug seeking for both substances and found that performance on the task was negatively correlated with WIN intake as well as WIN-seeking in abstinence. Worse performance on the working memory task was not significantly correlated with increased cocaine intake, but it was significantly correlated with increased cocaine-seeking during abstinence at certain delay lengths. Thus, while behavioral training that enhanced expression of BDNF in the prefrontal cortex did not provide a protective effect against drug taking or drug seeking, it did predict the extent of drug seeking or drug taking in individual animals. These results suggest that pre-existing differences in cognitive abilities may predict a particular vulnerability to drug taking and relapse-like behaviors after substance exposure.

Using our rodent models of drug self-administration, we questioned if cognitive training could have a preventative effect on substance use. Cognitive training has been proposed as an addition to existing psychotherapeutic treatments, yet incorporation of these methods has not reliably improved treatment outcomes across multiple psychiatric disorders, including substance use (Wanmaker et al., 2018; Redick, 2019; Gobet and Sala, 2023). Our results suggest that prior training is not effective in reducing substance use-associated behaviors and aligns with human studies indicating limited use for this type of cognitive training.

Despite this difference, use of preclinical rodent models has demonstrated that cognitive training methodologies can reduce cocaine-seeking (Boivin et al., 2015), and may still be useful for improving executive function across the lifespan (Stavroulaki et al., 2021). We found an interesting effect where individual differences in cognitive ability predicted substance use and potential for drug-seeking. Continued use of this model may help identify markers that predict vulnerability for development of a substance use disorder, or to predict responsiveness to treatment (Belin et al., 2016). While all animals in this study were adults at the time of training, individual differences in working memory performance during adolescence have been correlated with substance use in adulthood (Castellanos-Ryan et al., 2017; Morin et al., 2019). Evaluation and understanding of this relationship for individuals who may develop this, or other neuropsychiatric disorders could produce treatment strategies to improve outcomes in these specific individuals. Cognitive training may be uniquely useful in this subset of individuals, and this possibility warrants further study.

Our findings indicate that working memory training on both the simple and complex task was sufficient to increase BDNF protein concentration in the prelimbic prefrontal cortex and working memory task performance was mildly predictive of subsequent substance use, but this training was not sufficient to prevent drug-seeking. Rodent studies have indicated that BDNF measured in early adolescence along with working memory performance may be predictive of cocaine-seeking in male rats (Jordan and Andersen, 2018). In humans, the presence of genetic polymorphisms or measurements of BDNF levels in blood plasma are linked to improved outcomes after nonpharmacological treatment for neuropsychiatric disorders (Notaras et al., 2015). Little consensus has been reached about the magnitude of the beneficial link (Irwin et al., 2023). Prior studies utilizing exogenous BDNF administration into the medial prefrontal cortex have found consistent decreases in various types of reinstatement of cocaine seeking, suggesting that BDNF-induced molecular adaptations in the PFC-nucleus accumbens pathway are capable of reducing cocaine seeking (Berglind et al., 2009; Sun et al., 2015; Giannotti et al., 2018). One reason why we did not observe reduced drug-seeking from cognitive training induced increases in BDNF may be that a non-physiological enhancement of BDNF in this pathway is necessary to produce strong effects on drug-seeking behavior. Additionally, it may be that cognitive training did not increase BDNF if the neurons projecting to the nucleus accumbens. As the nucleus accumbens is highly involved in developing and maintaining drug-seeking behaviors, continued investigation into activating plasticity within these specific projections from the prefrontal cortex may lead to developing novel, non-invasive methods for promoting executive control over behavior and reducing drug-seeking.

This study focused on adult male rats and did not investigate any potential sex differences in the effect of cognitive training or BDNF expression. BDNF expression may differ between sexes, and some beneficial effects of BDNF may be more likely to emerge in males (Chan and Ye, 2017). The genetic polymorphism that promotes BDNF protein expression may be more likely to confer improved cognition after physical activity specifically in males (Watts et al., 2018), while the polymorphism that results in reduced BDNF expression is associated with psychiatric disorders such as major depression in men (Verhagen et al., 2010). Thus, the role of working memory training and BDNF signaling on drug seeking behaviors in females is not clear and warrants further investigation in future studies.

The interpretation of this study is limited by several factors including that this study utilized one unit dose of cocaine and WIN for all self-administration experiments, though these doses were chosen based on their known ability to promote reliable self-administration and reinstatement behavior (Rich et al., 2019; Bender and Torregrossa, 2021). The correlations between working memory performance and WIN self-administration or WIN-seeking in abstinence may suggest that this relationship only emerges for a relatively weak reinforcer, such as a cannabinoid, and not with a reinforcer that promotes resilient self-administration such as cocaine. Thus, it is possible that a stronger correlation between cognitive performance and drug intake could emerge at lower doses of cocaine. Additionally, this study utilized cognitive training prior to measurement of BDNF protein or drug exposure, which allowed us to identify protein levels and correlate behavioral performance in the absence of confounds related to drug exposure. However, this method did not directly model the experience of humans with substance use. Cognitive training is proposed as a component of treatment for substance use disorders as a method of recovering cognitive function lost due to drug use (Nixon and Lewis, 2019; Caetano et al., 2021). To address some of these questions, future studies can build on these results by directly associating endogenous BDNF after working memory training with drug seeking and working memory performance. Further, training paradigms that include punishment or incentives for inhibiting drug seeking can be incorporated into the methodology to more closely model human conditions, and measure how cognitive training may promote improved behavioral outcomes under those circumstances.

### Conclusion

While working memory training does produce changes in protein expression consistent with quelling drug-seeking behaviors, we found that this training was not sufficient to reduce addiction-associated behaviors of drug-taking or drug-seeking in abstinence. This is consistent with many recent findings indicating that cognitive training may not promote behavioral control that transfers to substance abuse or other psychiatric disorders. Working memory performance, however, may serve as a predictor for the magnitude of substance use.

## Acknowledgements

The authors would like to thank Megan Bertholomey, Jennifer Zeak, Brooke Bender, and Dana Smith for their outstanding technical expertise.

## Data Availability Statement

The data underlying this article will be shared on reasonable request to the corresponding author.

